# Training and inferring neural network function with multi-agent reinforcement learning

**DOI:** 10.1101/598086

**Authors:** Matthew Chalk, Gasper Tkacik, Olivier Marre

## Abstract

A central goal in systems neuroscience is to understand the functions performed by neural circuits. Previous top-down models addressed this question by comparing the behaviour of an ideal model circuit, optimised to perform a given function, with neural recordings. However, this requires guessing in advance what function is being performed, which may not be possible for many neural systems. To address this, we propose a new framework for optimising a recurrent network using multi-agent reinforcement learning (RL). In this framework, a reward function quantifies how desirable each state of the network is for performing a given function. Each neuron is treated as an ‘agent’, which optimises its responses so as to drive the network towards rewarded states. Three applications follow from this. First, one can use multi-agent RL algorithms to optimise a recurrent neural network to perform diverse functions (e.g. efficient sensory coding or motor control). Second, one could use inverse RL to infer the function of a recorded neural network from data. Third, the theory predicts how neural networks should adapt their dynamics to maintain the same function when the external environment or network structure changes. This could lead to theoretical predictions about how neural network dynamics adapt to deal with cell death and/or varying sensory stimulus statistics.

## Introduction

Neural circuits have evolved to perform a range of different functions, from sensory coding to muscle control and decision making. A central goal of systems neuroscience is to elucidate what these functions are and how neural circuits implement them. A common ‘top-down’ approach starts by formulating a hypothesis about the function performed by a given neural system (e.g. efficient coding/decision making), which can be formalised via an objective function [1–10]. This hypothesis is then tested by comparing the predicted behaviour of a model circuit that maximises the assumed objective function (possibly given constraints, such as noise/metabolic costs etc.) with recorded responses.

One of the earliest applications of this approach was sensory coding, where neural circuits are thought to efficiently encode sensory stimuli, with limited information loss [7–13]. Over the years, top-down models have also been proposed for many central functions performed by neural circuits, such as generating the complex patterns of activity necessary for initiating motor commands [3], detecting predictive features in the environment [4], or memory storage [5]. Nevertheless, it has remained difficult to make quantitative contact between top-down model predictions and data, in particular, to rigorously test which (if any) of the proposed functions is actually being carried out by a real neural circuit.

The first problem is that a pure top-down approach requires us to hypothesise the function performed by a given neural circuit, which is often not possible. Second, even if our hypothesis is correct, there may be multiple ways for a neural circuit to perform the same function, so that the predictions of the top-down model may not match the data.

Here we propose a new framework for considering optimal coding by a recurrent neural network, that aims to overcome these problems. First, we show how optimal coding by a recurrent neural network can be re-cast as a multi-agent reinforcement learning (RL) problem [14–18] (Fig 1). In this framework, a reward function quantifies how desirable each state of the network is for performing a given computation. Each neuron is then treated as a separate ‘agent’, which optimises its responses (i.e. when to fire a spike) so as to drive the network towards rewarded states, given a constraint on the information each neuron encodes about its inputs. This framework is very general – different choices of reward function result in the network performing diverse functions, from efficient coding to decision making and optimal control – and thus has the potential to unify many previous theories of neural coding.

**Fig 1.**
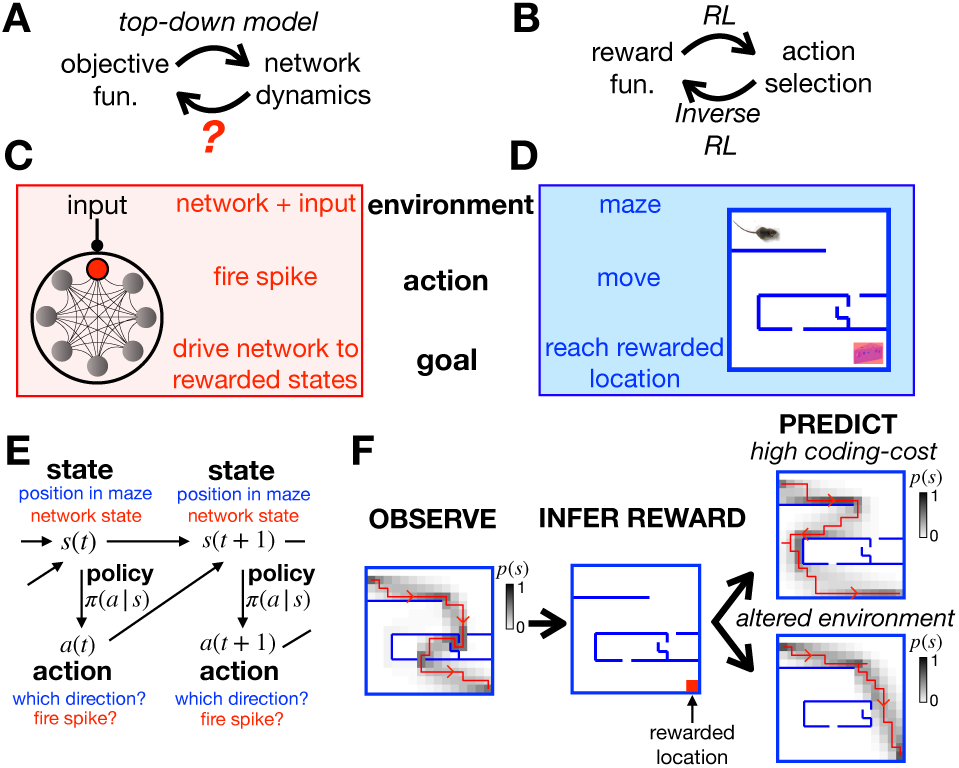
General approach. (**A**) Top-down models use an assumed objective function to derive the optimal neural dynamics. The inverse problem is to infer the objective function from observed neural responses. (**B**) RL uses an assumed reward function to derive an optimal set of actions that an agent should perform in a given environment. Inverse RL infers the reward function from the agent’s actions. (**C-D**) A mapping between the neural network and textbook RL setup. (**E**) Both problems can be formulated as MDPs, where an agent (or neuron) can choose which actions, *a*, to perform to alter their state, *s*, and increase their reward. (**F**) Given a reward function and coding cost (which penalises complex policies), we can use entropy-regularised RL to derive the optimal policy (left). Here we plot a single trajectory sampled from the optimal policy (red), as well as how often the agent visits each location (shaded). Conversely, we can use inverse RL to infer the reward function from the agent’s policy (centre). We can then use the inferred reward to predict how the agent’s policy will change when we increase the coding cost to favour simpler (but less rewarded) trajectories (top right), or move the walls of the maze (bottom right).

Next, we show how our proposed framework could be used to tackle the inverse problem, of inferring the reward function from the observed network dynamics. Previous work has proposed ‘inverse RL’ algorithms for inferring the original reward function from an agent’s actions [20–24]. Here we show how this framework can be adapted to infer the reward function optimised by a recurrent neural network. Further, given certain conditions we show that the reward function can be expressed as a closed-form expression of the observed network dynamics.

We hypothesise that the inferred reward function, rather than e.g. the properties of individual neurons, is the most succinct mathematical summary of the network, that generalises across different contexts and conditions. Thus we could use our framework to quantitatively predict how the network will adapt or learn in order to perform the same function when the external context (e.g. stimulus statistics), constraints (e.g. noise level) or the structure of the network (e.g. due to cell death or experimental manipulation) change. Our framework could thus not only allows RL to be used to train neural networks and use inverse RL to infer their function, but also could generate predictions for a wide range of experimental manipulations.

## Results

### General approach

We can quantify how well a network performs a specific function (e.g. sensory coding/decision making) via an objective function *L*_*π*_ (where *π* denotes the parameters that determine the network dynamics) (Fig 1A). There is a large literature describing how to optimise the dynamics of a neural network, *π*, to maximise specific objective functions, *L*_*π*_, given constraints (e.g. metabolic cost/wiring constraints etc.) [1–10]. However, it is generally much harder to go in the opposite direction, to infer the objective function, *L*_*π*_, from observations of the network dynamics.

To address this question, we looked to the field of reinforcement learning (RL) [14–18], which describes how an agent should choose actions so as to maximise the reward they receive from their environment (Fig 1B). Conversely, another paradigm, called inverse RL [20–24], explains how to go in the opposite direction, to infer the reward associated with different states of the environment from observations of the agent’s actions. We reasoned that, if we could establish a mapping between optimising neural network dynamics (Fig 1A) and optimising an agent’s actions via RL (Fig 1B), then we could use inverse RL to infer the objective function optimised by a neural network from its observed dynamics.

To illustrate this, let us compare the problem faced by a single neuron embedded within a recurrent neural network (Fig 1C) to the textbook RL problem of an agent navigating a maze (Fig 1D). The neuron’s environment is determined by the activity of other neurons in the network and its external input; the agent’s environment is determined by the walls of the maze. At each time, the neuron can choose whether to fire a spike, so as to drive the network towards states that are ‘desirable’ for performing a given function; at each time, the agent in the maze can choose which direction to move in, so as to reach ‘desirable’ locations, associated with a high reward.

Both problems can be formulated mathematically as Markov Decision Processes (MDPs) (Fig 1E). Each state of the system, *s* (i.e. the agent’s position in the maze, or the state of the network and external input), is associated with a reward, *r*(*s*). At each time, the agent can choose to perform an action, *a* (i.e. moving in a particular direction, or firing a spike), so as to reach a new state *s*′ with probability, *p*(*s*′|*a, s*). The probability that the agent performs a given action in each state, *π*(*a*|*s*), is called their policy.

We assume that the agent (or neuron) optimises their policy to maximise their average reward, 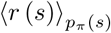 (where 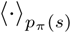 denotes the average over the steady state distribution, *p*_*π*_ (*s*), with a policy *π* (*a*|*s*)), given a constraint on the information they can encode about their state, *I*_*π*_(*a*; *s*) (this corresponds, for example, to constraining how much a neuron can encode about the rest of the network and external input). This can be achieved by maximising the following objective function:

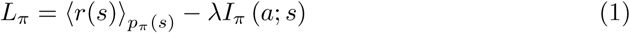

where *λ* is a constant that controls the strength of the constraint. Note that in the special case where the agent’s state does not depend on previous actions (i.e. *p* (*s*′|*a, s*) = *p* (*s*′*s*)) and the reward depends on their current state and action, this is the same as the objective function used in rate-distortion theory [25, 26]. We can also write the objective function as:

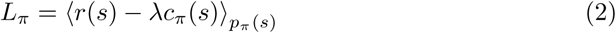

where *c*_*π*_(*s*) is a ‘coding cost’, equal to the Kullback-Leibler divergence between the agent’s policy and the steady-state distribution over actions, *D*_*KL*_ [*π* (*a*|*s*)‖*p*_*π*_ (*a*)]. We hereon refer to the difference, *r*(*s*) − *λc*_*π*_(*s*), as the ‘return’ associated with each state.

In Methods section we show how this objective function can be maximised via entropy-regularised RL [15, 18] to obtain the optimal policy, which satisfies the relation:

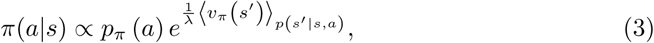

where *v*_*π*_ (*s*) is the ‘value’ associated with each state, defined as the total return predicted in the future if the agent starts in a given state, minus the average return, *L*_*π*_:

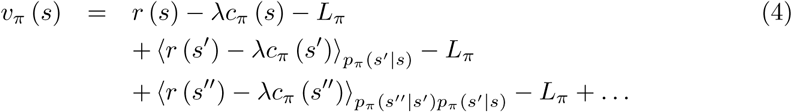

where *s, s*′ and *s*″ denote three consecutive states of the agent. Subtracting the average return, *L*_*π*_, from each term in the sum ensures that this series converges to a finite value [19]. Thus, actions that drive the agent towards high-value states are preferred over actions that drive the agent towards low value states. Note the difference between a state’s value, *v* (*s*), and its return, *r* (*s*) − *λc*_*π*_ (*s*): a state with low return can nonetheless have a high-value if it allows the agent to transition to other states associated with a high return in the future.

Let us return to our toy example of the agent in a maze. Figure 1F (left) shows the agent’s trajectory through the maze after optimising their policy using entropy regularized RL to maximise *L*_*π*_ (Methods, section). In this example, a single location, in the lower-right corner of the maze, has a non-zero reward (Fig 1F, centre). However, suppose we didn’t know this; could we infer the reward at each location just by observing the agent’s trajectory in the maze? In Methods section we show that this can be done by finding the reward function that maximises the log-likelihood of the optimal policy, averaged over observed actions and states, ⟨log *π*\* (*a*|*s*) ⟩_data_. If the coding cost is non-zero (*λ* > 0), this problem is generally well-posed, meaning there is a unique solution for *r*(*s*).

Once we have inferred the reward function optimised by the agent, we can then use it to predict how their behaviour will change when we alter their external environment or internal constraints. For example, we can predict how the agent’s trajectory through the maze will change when we move the position of the walls (Fig 1F, lower right), or increase the coding cost so as to favour simpler (but less rewarded) trajectories (Fig 1F, upper right).

### Optimising neural network dynamics

We used these principles to infer the function performed by a recurrent neural network. We considered a model network of *n* neurons, each described by a binary variable, *σ*_*i*_ = −1*/*1, denoting whether the neuron is silent or spiking respectively (Methods section). The network receives an external input, ***x***. The network state is described by an *n*-dimensional vector of binary values, ***σ*** = (*σ*_1_, *σ*_2_, …, *σ*_*n*_)^*T*^. Both the network and external input have Markov dynamics. Neurons are updated asynchronously: at each time-step a neuron is selected at random, and its state updated by sampling from 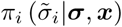. The dynamics of the network are fully specified by the set of response probabilities, 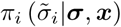, and input statistics, *p*(***x***′ ***x***).

As before, we use a reward function, *r* (***σ, x***), to express how desirable each state of the network is to perform a given functional objective. For example, if the objective of the network is to faithfully encode the external input, then an appropriate reward function might be the negative squared error: 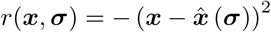, where 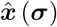 denotes an estimate of ***x***, inferred from the network state, ***σ***. More generally, different choices of reward function can be used to describe a large range of functions that may be performed by the network.

The dynamics of the network, *π*, are said to be optimal if they maximise the average reward, 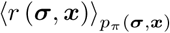, given a constraint on the information each neuron encodes about the rest of the network and external inputs, 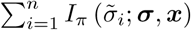. This corresponds to maximising the objective function:

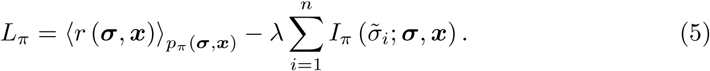

where *λ* controls the strength of the constraint. For each neuron, we can frame this optimisation problem as an MDP, where the state, action, and policy correspond to the network state and external input {***σ, x***}, the neuron’s proposed update 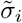, and the response probability 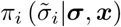, respectively. Thus, we can optimise the network dynamics, by treating each neuron as an agent, and optimising its response probability, 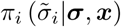, via entropy-regularised RL, as we did for the agent in the maze. Further, as each update increases the objective function *L*_*π*_, we can alternate updates for different neurons to optimise the dynamics of the entire network (as in multi-agent RL). In Methods section, we show that this results in optimal response probabilities that satisfy the relation:

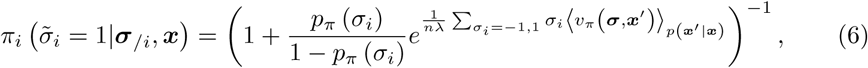

where ***σ***_*/i*_ denotes the state of all neurons except for neuron *i*, and *v*_*π*_ (***σ, x***) is the value associated with each state:

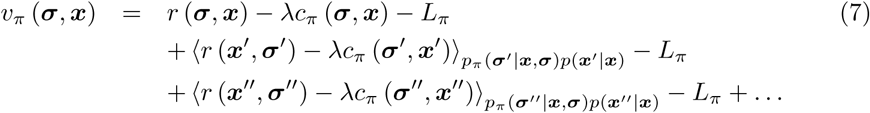

The coding cost, 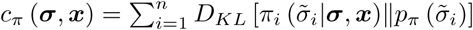 penalises deviations from each neuron’s average firing rate. The network dynamics are optimised by alternately updating the value function and neural response probabilities until convergence (Methods section).

To see how this works in practice, we simulated a network of 8 neurons that receive a binary input *x* (Fig 2A). The assumed goal of the network is to fire exactly 2 spikes when *x* = −1, and 6 spikes when *x* = 1, while minimising the coding cost. To achieve this, the reward was set to unity when the network fired the desired number of spikes, and zero otherwise (Fig 2B). Using entropy-regularised RL, we derived optimal tuning curves for each neuron, which show how their spiking probability should optimally vary depending on the input, *x*, and number of spikes fired by other neurons (Fig 2C). We confirmed that after optimisation the number of spikes fired by the network was tightly peaked around the target values (Fig 2D). Decreasing the coding cost reduced noise in the network, decreasing variability in the total spike count.

**Fig 2.**
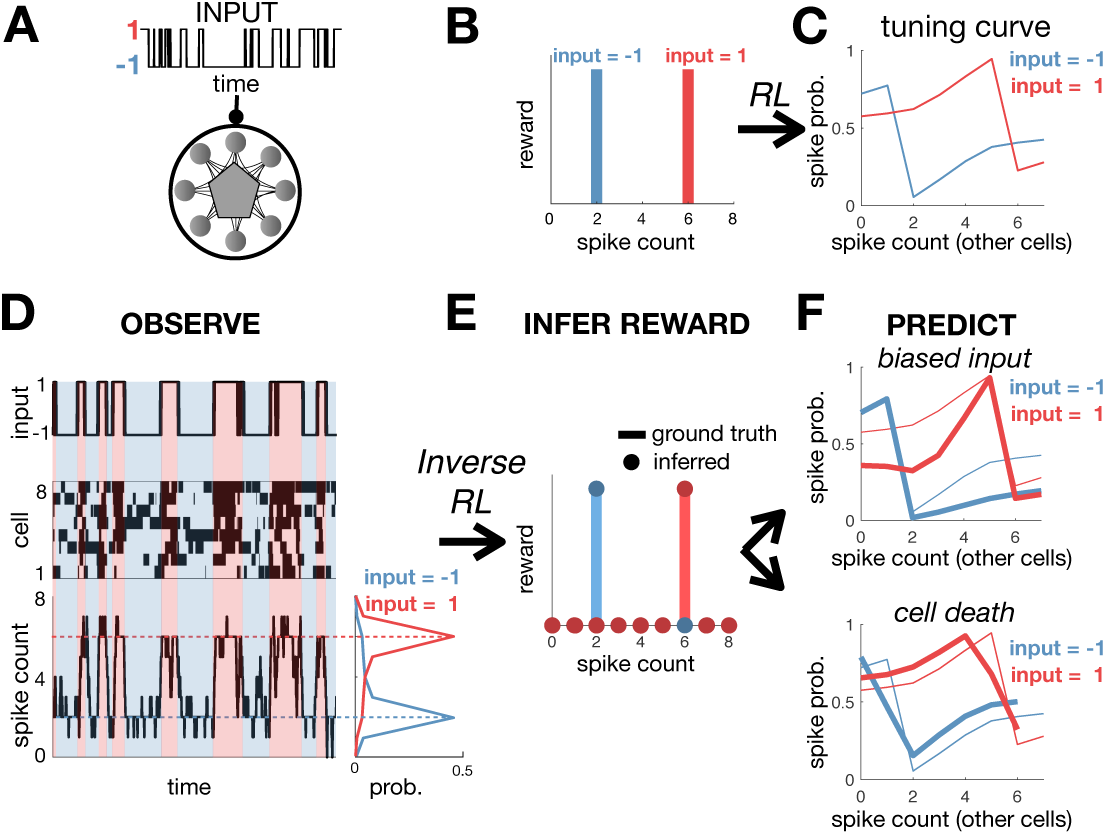
Training and inferring the function performed by a neural network. (**A**) A recurrent neural network receives a binary input, *x*. (**B**) The reward function equals 1 if the network fires 2 spikes when *x* = −1, or 6 spikes when *x* = 1. (**C**) After optimisation, neural tuning curves depend on the input, *x*, and total spike count. (**D**) Simulated dynamics of the network with 8 neurons (left). The total spike count (below) is tightly peaked around the rewarded values. (**E**) Using inverse RL on the observed network dynamics, we infer the original reward function used to optimise the network from its observed dynamics. (**F**) The inferred reward function is used to predict how neural tuning curves will adapt depending on contextual changes, such as varying the input statistics (e.g. decreasing *p*(*x* = 1)) (top right), or cell death (bottom right). Thick/thin lines show adapted/original tuning curves, respectively.

### Inferring the objective function from the neural dynamics

We next asked if we could use inverse RL to infer the reward function optimised by a neural network, just from its observed dynamics (Fig 2D). For simplicity, let us first consider a recurrent network that receives no external input. In this case, the optimal dynamics (Eqn 6) correspond to Gibbs sampling from a steady-state distribution: 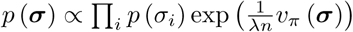. We can combine this with the Bellmann equality, which relates the reward, value and cost functions (according to: *r* (***σ***) = *v*_*π*_ (***σ***) + *λc*_*π*_ (***σ***) − ⟨*v*_*π*_ (***σ***′)⟩_*p*(***σ′*|*σ***)_ + *L*_*π*_; see Methods) to derive an expression for the reward function:

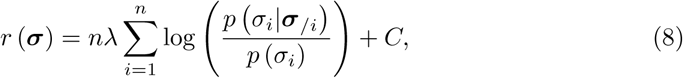

where *p* (*σ*_*i*_|***σ***_*/i*_) denotes the probability that neuron *i* is in state *σ*_*i*_, given the current state of all the other neurons and *C* is an irrelevant constant (see Methods). Without loss of generality, we can set the coding cost, *λ*, to 1 (since altering *λ* rescales the inferred reward and coding cost by the same factor, rescaling the objective function without changing its shape). In the Methods, we show how we can recover the reward function when there is an external input. In this case, we do not obtain a closed-form expression for the reward function, but must instead infer it via maximum likelihood.

Figure 2E shows how we can use inverse RL to infer the reward function optimised by a model network from its observed dynamics, in the presence of an external input. Note, that our method did not make any *a priori* assumptions about the parametric form of the reward function, which was allowed to vary freely as a function of the network state and input, (***σ, x***). Nonetheless, we can use a simple clustering algorithm (e.g. k-means) to recover the fact that the inferred reward took two binary values; further analysis reveals that the reward is only non-zero when the network fired exactly 2 spikes when *x* = −1, and 6 spikes when *x* = 1. As for the agent in the maze, we can use this inferred reward function to predict how the network dynamics will vary depending on the internal/external constraints. For example, we can predict how neural tuning curves will vary if we alter the input statistics (Fig 2F, upper), or remove a cell from the network (Fig 2F, lower).

Our ability to correctly infer the reward function optimised by the network will be fundamentally limited by the amount of available data. Fig 3A shows how the correlation between the inferred and true reward increases with the amount of data samples used to infer the reward. (Note that each discrete time-step is considered to be one data sample.) As the number of samples is increased, the distribution of inferred rewards becomes more tightly peaked around two values (Fig 3B), reflecting the fact that the true reward function was binary. Of course, with real neural data we will not have access to the ‘true’ reward function. In this case, we can test how well our inferred reward function is able to predict neural responses in different conditions. Figure 3C-D shows how the predicted response distribution (when we alter the input statistics, Fig 3C, or remove cells, Fig 3D) becomes more accurate as we increase the number of samples used to estimate the reward function.

**Fig 3.**
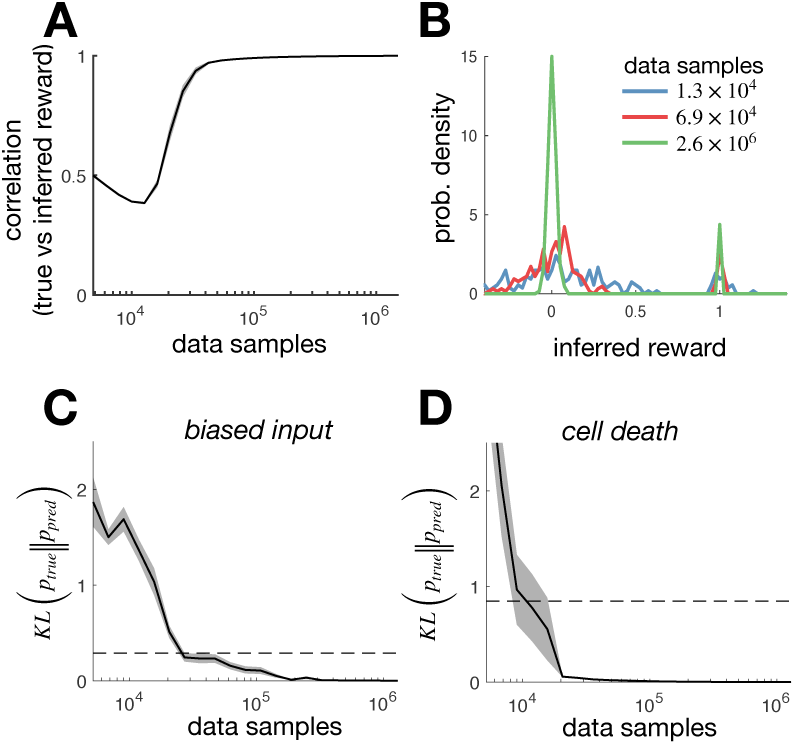
Inferring the reward from limited data. (**A**) The *r*^2^-goodness of fit between the true reward, and the reward inferred using a finite number of samples (a sample is defined as an observation of the network state at a single time-point). The solid line indicates the *r*^2^ value averaged over 20 different simulations, while the shaded areas indicate the standard error on the mean. (**B**) Distribution of rewards inferred from a variable numbers of data samples. As the number of data samples is increased, the distribution of inferred rewards becomes more sharply peaked around 0 and 1 (reflecting the fact that the true reward was binary). (**C**) The KL-divergence between the optimal response distribution with altered input statistics (see Fig 2F, upper) and the response distribution predicted using the reward inferred in the initial condition from a variable number of samples. The solid line indicates the KL-divergence averaged over 20 different simulations, while the shaded areas indicate the standard error on the mean. A horizontal dashed line indicates the KL-divergence between the response distribution with biased input and the original condition (that was used to infer the reward). (**D**) Same as panel (C), but where instead of altering the input statistics, we remove cells from the network (see Fig 2F, lower).

### Inferring efficiently encoded stimulus features

An influential hypothesis, called ‘efficient coding’, posits that sensory neural circuits have evolved to encode maximal information about sensory stimuli, given internal constraints [7–13]. However, the theory does not specify which stimulus features are relevant to the organism, and thus should be encoded. Here we show how one could use inverse RL to: (i) infer which stimulus features are encoded by a recorded neural network, and (ii) test whether these features are encoded efficiently.

Efficient coding posits that neurons maximise information encoded about some relevant feature, ***y*** (***x***), given constraints on the information encoded by each neuron about their inputs, ***x*** (Fig 4A). This corresponds to maximising:

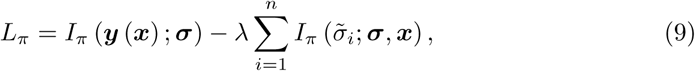

where *λ* controls the strength of the constraint. Noting that the second term is equal to the coding cost we used previously (Eqn 5), we can rewrite this objective function as:

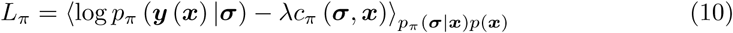

where we have omitted terms which don’t depend on *π*. Now this is exactly the same as the objective function we have been using so far (Eqn 5), in the special case where the reward function, *r* (***σ, x***), is equal to the log-posterior, log *p*_*π*_ (***y*** (***x***)|***σ***). As a result we can maximise *L*_*π*_ via an iterative algorithm, where on each iteration we update the reward function by setting *r* (***x, σ***) ←log *p*_*π*_(***y*** (***x***) |***σ***), before then optimising the network dynamics, via entropy-regularised RL. Thus, thanks to the correspondence between entropy-regularised RL and efficient coding we could derive an algorithm to optimise the dynamics of a recurrent network to perform efficient coding [28].

**Fig 4.**
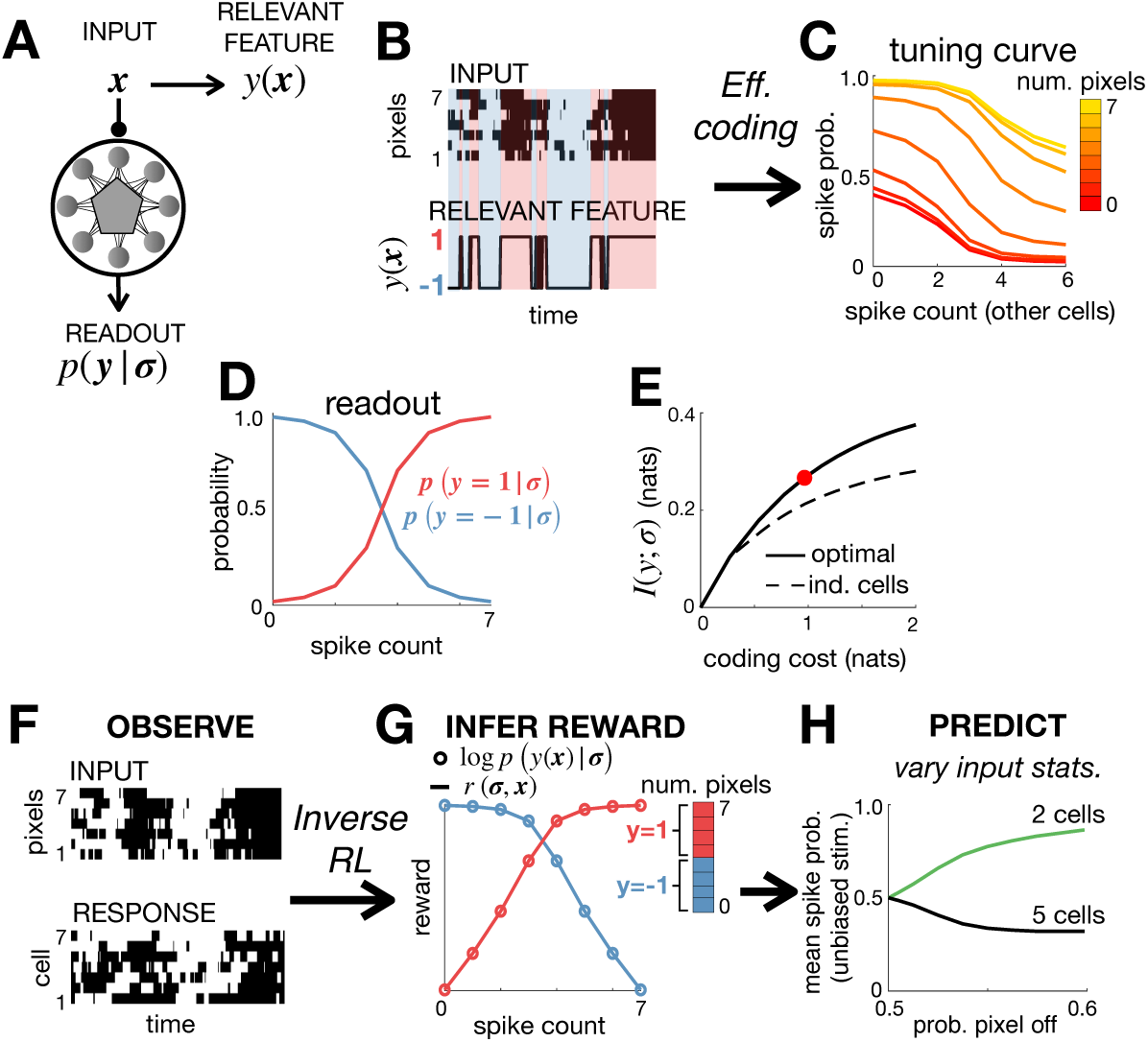
Efficient coding and inverse RL. (**A**) The neural code was optimised to efficiently encode an external input, ***x***, so as to maximise information about a relevant stimulus feature *y* (***x***). (**B**) The input, ***x*** consisted of 7 binary pixels. The relevant feature, *y* (***x***), was equal to 1 if >3x pixels were active, and −1 otherwise. (**C**) Optimising a network of 7 neurons to efficiently encode *y* (***x***) resulted in all neurons having identical tuning curves, which depended on the number of active pixels and total spike count. (**D**) The posterior probability that *y* = 1 varied monotonically with the spike count. (**E**) The optimised network encoded significantly more information about *y* (***x***) than a network of independent neurons with matching stimulus-dependent spiking probabilities, *p* (*σ*_*i*_ = 1|***x***). The coding cost used for the simulations in the other panels is indicated by a red circle. (**F**-**G**) We use the observed responses of the network (**F**) to infer the reward function optimised by the network, *r* (***σ, x***) (**G**). If the network efficiently encodes a relevant feature, *y* (***x***), then the inferred reward (solid lines) should be proportional to the log-posterior, log *p* (*y* (***x***) |***σ***) (empty circles). This allows us to (i) recover *y* (***x***) from observed neural responses, (ii) test whether this feature is encoded efficiently by the network. (**H**) We can use the inferred objective to predict how varying the input statistics, by reducing the probability that pixels are active, causes the population to split into two cell types, with different tuning curves and mean firing rates (right).

As an illustration, we simulated a network of 7 neurons that receive a sensory input consisting of 7 binary pixels (Fig 4B, top). In this example, the ‘relevant feature’, *y* (***x***) was a single binary variable, which was equal to 1 if 4 or more pixels were active, and −1 otherwise (Fig 4B, bottom). Using the efficient-coding algorithm described above, we derived optimal tuning curves, showing how each neuron’s spiking probability should vary with both the number of active pixels and number of spikes fired by other neurons (Fig 4C). We also derived how the optimal readout, *p* (*y*|***σ***), should depend on the number of spiking neurons (Fig 4D). Finally, we verified that the optimised network encodes significantly more information about the relevant feature than a network of independent neurons, over a large range of coding costs (Fig 4E).

Now, imagine that we just observe the stimulus and neural responses (Fig 4F). Can we recover the relevant feature, *y* (***x***)? To do this, we first use inverse RL to infer the reward function from observed neural responses (in exactly the same way as described in the previous section) (Fig 4G). As before, we made no *a priori* assumptions about the parametric form of the reward function, which was allowed to vary freely as a function of the network state and input, (***σ, x***). Now, as described above, if the network is performing efficient coding then the inferred reward, *r* (***σ, x***) should be proportional to the log-posterior, log *p* (*y* (***x***)|***σ***). Thus, given ***σ***, the inferred reward, *r* (***σ, x***) should only depend on changes to the input, ***x***, that alter *y* (***x***). As a result, we can use the inferred reward to uncover all inputs, ***x***, that map onto the same value of *y* (***x***). In our example, we see that the inferred reward collapses onto two curves only (blue and red in Fig 4G), depending on the total number of pixels in the stimulus. This allows us to deduce that the relevant coded variable, *y* (***x***), must be a sharp threshold on the number of simultaneously active pixels. In contrast, the neural tuning curves vary smoothly with the number of active pixels (Fig 4C). Next, having recovered *y* (***x***), we can check whether it is encoded efficiently by seeing whether the inferred reward, *r* (***σ, x***) is proportional to the log-posterior, log *p* (*y* (***x***)|***σ***).

Note that our general approach could also generalise to more complex efficient coding models, where the encoded variable, *y*, is not a binary function of the input, ***x***. In this case, we can perform a cluster analysis (e.g. k-means) to reveal which inputs, ***x***, map onto similar reward. If the network is performing efficient coding then these inputs should also map onto the same encoded feature, *y* (***x***).

Finally, once we have inferred the function performed by the network, we can predict how its dynamics will vary with context, such as when we alter the input statistics. For example, in our simulation, reducing the probability that input pixels are active causes the neural population to split into two cell-types, with distinct tuning curves and mean firing rates (Fig 4H) [13].

### Parametric model of neural responses

The basic framework described above is limited by the fact that the number of states, *n*_*s*_, scales exponentially with the number of neurons (*n*_*s*_ = 2^*n*^). Thus, it will quickly become infeasible to compute the optimal dynamics as the number of neurons increases. Likewise, we will need an exponential amount of data to reliably estimate the sufficient statistics of the network, required to infer the reward function (Fig 3).

For larger networks, this problem can be circumvented by using tractable parametric approximation of the value function and reward functions. As an illustration, let us consider a network with no external input. If we approximate the value function by a quadratic function of the responses, our framework predicts a steady-state response distribution of the form: 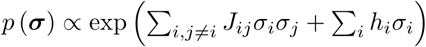, where *J*_*ij*_ denotes the pairwise couplings between neurons, and *h*_*i*_ is the bias. This corresponds to a pairwise Ising model, which has been used previously to model recorded neural responses [27, 28]. (Note that different value functions could be used to give different neural models; e.g. choosing *v* (***x***) = *f* (***w* ·*x***), where ***x*** is the feed-forward input, results in a linear-nonlinear neural model.) In Methods section we derive an algorithm to optimise the coupling matrix, ***J***, for a given reward function and coding cost.

To illustrate this, we simulated a network of 12 neurons arranged in a ring, with reward function equal to 1 if exactly 4 adjacent neurons are active together, and 0 otherwise. After optimisation, nearby neurons were found to have positive couplings, while distant neurons had negative couplings (Fig 5A). The network dynamics generate a single hill of activity which drifts smoothly in time. This is reminiscent of ring attractor models, which have been influential in modeling neural functions such as the rodent and fly head direction system [29–31]. (Indeed, eqn. 6 suggests why our framework generally leads to attractor dynamics, as each transition will tend to drive the network to higher-value ‘attractor states’.)

**Fig 5.**
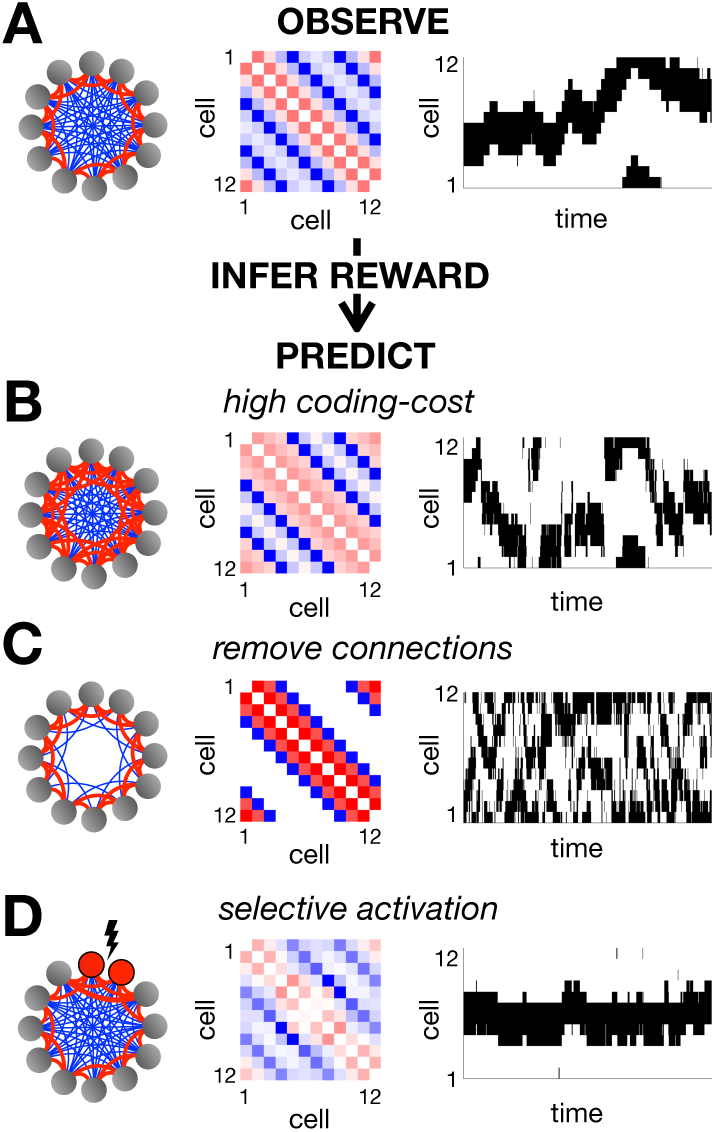
Pairwise coupled network. (**A**) We optimized the parameters of a pairwise coupled network, using a reward function that was equal to 1 when exactly 4 adjacent neurons were simultaneously active, and 0 otherwise. The resulting couplings between neurons are schematized on the left, with positive couplings in red and negative couplings in blue. The exact coupling strengths are plotted in the centre. On the right we show an example of the network dynamics. Using inverse RL, we can infer the original reward function used to optimise the network from its observed dynamics. We can then use this inferred reward to predict how the network dynamics will vary when we increase the coding cost (**B**), remove connections between distant neurons (**C**) or selectively activate certain neurons (**D**).

As before, we can then use inverse RL to infer the reward function from the observed network dynamics. However, note that when we use a parametric approximation of the value function this problem is not well-posed, and we have to make additional assumptions about the form of the reward function. We first assumed a ‘sparse’ reward function, where only a small number of states, ***σ***, are assumed to be associated with non-zero positive reward (see Methods). Using this assumption, we could well recover the true reward function from observations of the optimised neural responses (with an *r*^2^ value greater than 0.9).

Having inferred the reward function optimised by the network, we can then use it to predict how the coupling matrix, ***J***, and network dynamics will vary if we alter the internal/external constraints. For example, we can use the inferred reward to predict how increasing the coding cost will result in stronger positive couplings between nearby neurons and a hill of activity that sometimes jumps discontinuously between locations (Fig 5B); removing connections between distant neurons will result in two uncoordinated peaks of activity (Fig 5C); finally, selectively activating certain neurons will ‘pin’ the hill of activity to a single location (Fig 5D).

To illustrate the effect of assuming different reward functions, we considered two different sets of assumptions (in addition to the sparse model, described above): a ‘pairwise model’, where the reward is assumed to be a quadratic function of the network state, and a ‘global model’ where the reward is assumed to depend only on global spike count (see Methods). In all three cases, the inferred reward function provided a reasonable fit to the true reward function, (averaged over states visited by the network; Fig 6A). However, only the sparse and pairwise models were able to predict how neural responses changed when, for example, we optimised the network with a higher coding-cost (Fig 6B).

**Fig 6.**
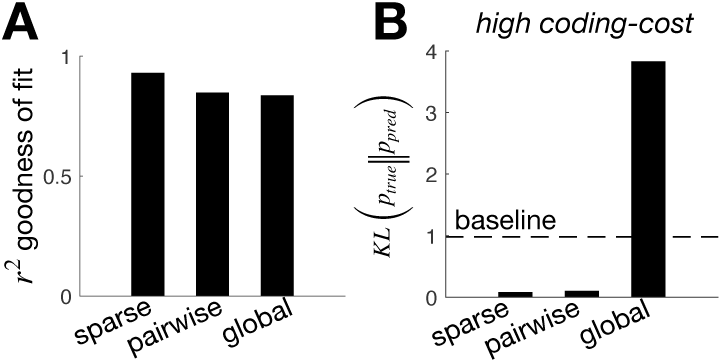
Effect of assuming different types of reward function. We compared the inferred reward when we assumed a sparse model (i.e. a small number of states associated with non-zero positive reward) a pairwise model (i.e. the reward depends on the first and second-order response statistics) and a global model (i.e. the reward depends on the total number of active neurons only). (**A**) *r*^2^ goodness of fit between the true and the inferred reward, assuming a sparse, pairwise, or global model. (**B**) The KL-divergence between the optimal response distribution with high coding cost (see Fig 5B) and the response distribution predicted using the reward inferred in the initial condition, assuming a sparse, pairwise, or global model. A horizontal dashed line indicates the KL-divergence between the response distribution with high-coding cost and the original condition (that was used to infer the reward).

## Discussion

A large research effort has been devoted to developing ‘top-down’ models, which describe the network dynamics required to optimally perform a given function (e.g. decision making [6], control [3], efficient sensory coding [8] etc.). Here we show that optimising a recurrent neural network can often be formulated as a multi-agent RL problem. This insight can provide new ways to: (i) train recurrent networks to perform a given function; (ii) infer the function optimised by a network from recorded data; (iii) predict the dynamics of a network upon perturbation.

An alternative ‘bottom-up’ approach is to construct phenomenological models describing how neurons respond to given sensory stimuli and/or neural inputs [27, 28, 32, 33]. In common with our work, such models are directly fitted to neural data. However, these models generally do not set out to reveal the function performed by the network. Further, they are often poor at predicting neural responses in different contexts (e.g. varying stimulus statistics). Here we hypothesize that it is the function performed by a neural circuit that remains invariant, not its dynamics or individual cell properties. Thus, if we can infer what this function is, we should be able to predict how the network dynamics will adapt depending on the context, so as to perform the same function under different constraints. As a result, our theory could predict how the dynamics of a recorded network will adapt in response to a large range of experimental manipulations, such as varying the stimulus statistics, blocking connections, knocking out/stimulating cells etc. (Note however, that to predict how the network adapts to these changes, we will need to infer the ‘full’ reward function; in some cases, this may require measuring neural responses in multiple environments.)

Another approach is to use statistical methods, such as dimensionality reduction techniques, to infer information about the network dynamics (such as which states are most frequently visited), which can then be used to try and gain insight about the function it performs [34–37]. For example, in the context of sensory coding, an approach called ‘maximally informative dimensions’ seeks to find a low-dimensional projection of the stimulus that is most informative about a neuron’s responses [38]. However, in contrast to our approach, such approaches do not allow us to recover the objective function optimised by the network. As a result, they do not predict how neural responses will alter depending on the the internal/external constraints. It is also not clear how to relate existing dimensionality reduction methods, such as PCA, to the dynamics of a *recurrent* neural network (e.g. are certain states visited frequently because of the reward, external stimulus, or internal dynamics?). Nonetheless, in future work it could be interesting to see if dimensionality reduction techniques could be used to first recover a compressed version of the data, from which we could more easily use inverse RL methods to infer the objective function.

There is an extensive literature on how neural networks could perform RL [39–41]. Our focus here was different: we sought to use tools from RL and inverse RL to infer the function performed by a recurrent neural network. Thus, we do not assume the network receives an explicit reward signal: the reward function is simply a way of expressing which states of the network are useful for performing a given function. In contrast to previous work, we treat each neuron as an independent agent, which optimises their responses to maximise the reward achieved by the network, given a constraint on how much they can encode about their inputs. As well as being required for biological realism, the coding constraint has the benefit of making the inverse RL problem well-posed. Indeed, under certain assumptions, we show that it is possible to write a closed form expression for the reward function optimised by the network, given its steady-state distribution (Eqn 40).

Our framework relies on several assumptions about the network dynamics. First, we assume that the network has Markov dynamics, such that its state depends only on the preceding time-step. To relax this assumption, we could redefine the network state to include spiking activity in several time-steps. For example, we could thus include the fact that neurons are unlikely to fire two spikes within a given temporal window, called their refractory period. Of course, this increase in complexity would come at the expense of decreased computational tractability, which may necessitate approximations. Second, we assume the only constraint that neurons face is a ‘coding cost’, which limits how much information they encode about other neurons and external inputs. In reality, biological networks face many other constraints, such as the metabolic cost of spiking and constraints on the wiring between cells. Some of these constraints could be included explicitly as part of the inference process. For example, by assuming a specific form of approximate value function (e.g. a quadratic function of the neural responses), we can include explicit constraints about the network connectivity (e.g. pairwise connections between neurons). Other constraints (e.g. the metabolic cost of firing a spike), that are not assumed explicitly can be incorporated implicitly as part of the inferred reward function (e.g. a lower inferred reward for metabolically costly states, associated with high firing rates).

In addition to assumptions about the network dynamics, for large networks we will also need to assume a particular parametric form for the reward function. For example, in the context of efficient coding (Fig 4), this is equivalent to assuming a particular form for the decoder model (e.g. a linear decoder) that ‘reads-out’ the encoded variable from neural responses [10]. To test the validity of such assumptions, we could see how well the reward function, inferred in one context, was able to generalise to predict neural responses in other contexts.

For our framework to make predictions, the network must adapt its dynamics to perform the same function under different internal/external constraints. Previous work suggests that this may hold in certain cases, in response to changes in stimulus statistics [42, 43], or optogonetic perturbations [44]. Even so, it may be that real neural networks only partially adapt to their new environments. It would thus be interesting, in the future, to extend our framework to deal with this. For example, we could assess whether contextual changes in the neural responses enable the network to ascend the gradient of the objective function (given the new constraints) as predicted by our model.

A central tenet of our work is that the neural network has evolved to perform a specific function optimally, given constraints. As such, we could obtain misleading results if the recorded network is only approximately optimal. To deal with this, recent work by one of the present authors [45] proposed a framework in which the neural network is assumed to satisfy a *known* optimality criterion *approximately*. In this work, the optimality criterion is formulated as a Bayesian prior, which serves to nudge the network towards desirable solutions. This contrasts with the work presented here, where the network is assumed to satisfy an *unknown* optimality criterion *exactly*. Future work could explore the intersection between these two approaches, where the network is assumed to perform an *unknown* optimality criterion *approximately*. In this case, one will likely need to limit the space of possible reward functions, so that inference problem remains well-posed.

Our work unifies several influential theories of neural coding, that were considered separately in previous work. For example, we show a direct link between entropy-regularised RL [15–18] (Fig 1), ring-attractor networks [29–31] (Fig 5), and efficient sensory coding [7–13] (Fig 4). Further, given a static network without dynamics, our framework is directly equivalent to rate-distortion theory [25, 26]. Many of these connections are non-trivial. For example, the problem of how to train a network to efficiently code its inputs remains an open avenue of research. Thus, the realisation that efficient sensory coding by a recurrent network can be formulated as a multi-agent RL problem could help develop of future algorithms for learning efficient sensory representations (e.g., in contrast with brute force numerical optimization as in [9]). Indeed, recent work has shown how, by treating a feed-forward neural network as a multi-agent RL system, one can efficiently train the network to perform certain using only local learning rules [46]. However, while interesting in its own right, the generality of our framework means that we could potentially apply our theory to infer the function performed by diverse neural circuits, that have evolved to perform a broad range of different functional objectives. This contrasts with previous work, where neural data is often used to test a single top-down hypothesis, formulated in advance.

## Methods

### Entropy-regularised RL

We consider a Markov Decision Process (MDP) where each state of the agent *s*, is associated with a reward, *r*(*s*). At each time, the agent performs an action, *a*, sampled from a probability distribution, *π*(*a*|*s*), called their policy. A new state, *s*′, then occurs with a probability, *p*(*s*′ |*s, a*).

We seek a policy, *π* (*a*|*s*), that maximises the average reward, constrained on the mutual information between between actions and states. This corresponds to maximising the Lagrangian:

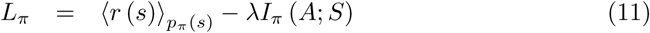

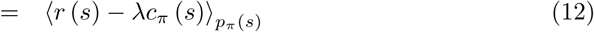

where *λ* is a lagrange-multiplier that determines the strength of the constraint, and *c*_*π*_ (*s*) = *D*_*KL*_ [*π* (*a*|*s*)‖*p*(*a*)].

Note that while in the rest of the paper we consider continuous tasks, where the Lagrangian is obtained by averaging over the steady-state distribution, our framework can also be applied without little changes to finite tasks, which occur within a time window, and where the Lagrangian is given by:

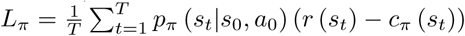. Unlike the continuous task described above, the optimal network dynamics may not have a corresponding equilibrium distribution [14].

Now, let us can define a value function:

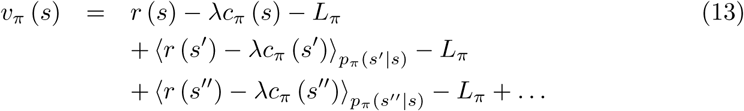

where *s, s*′ and *s*″ denote the agent’s state in three consecutive time-steps. We can write the following Bellmann equality for the value function:

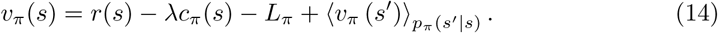

Now, consider the following greedy update of the policy, *π* (*a*|*s*):

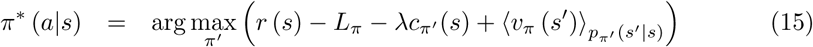

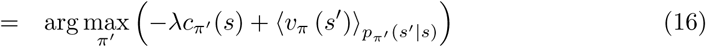

To preform this maximisation, we write the following Lagrangian:

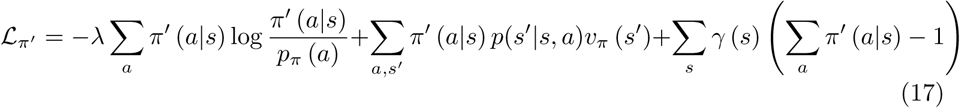

where the last term is required to enforce the constraint that ∑_*a*_ *π* (*a*|*s*) = 1. Setting the derivative to zero with respect to *γ* (*s*) and *π* (*a*|*s*), we have:

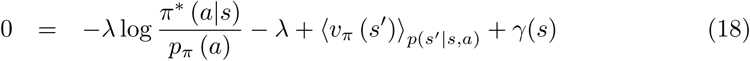

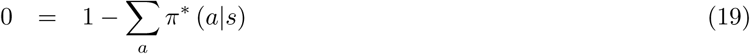

We can solve these equations to obtain the optimal greedy update:

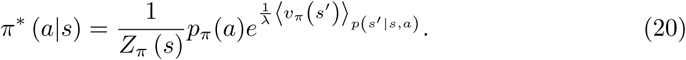

where *Z*_*π*_ (*s*) is a normalisation constant.

Using the policy improvement theorem [14], we can show that this greedy policy update is guaranteed to increase *L*_*π*_. To see this, we can substitute *π*\* into the Bellmann equation to give the following inequality:

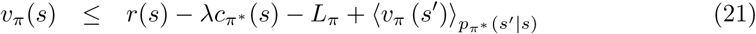

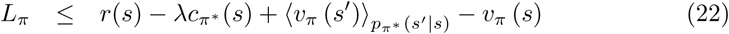

Next, we take the average of both sides with respect to the steady-state distribution, *p*_*π*\*_ (*s*):

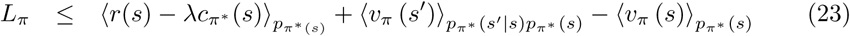

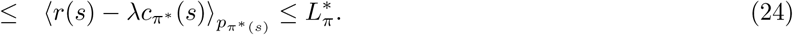

Thus, repeated application of the Bellmann recursion (Eqn 14) and greedy policy update (Eqn 20) will return the optimal policy, *π*\*(*a*|*s*), which maximises *L*_*π*_.

#### Inverse entropy-regularized RL

We can write the Bellmann recursion in Eqn 14 in vector form:

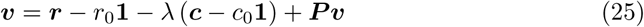

where ***v, c*** and ***r*** are vectors with elements, *v*_*s*_ ≡ *v* (*s*), *c*_*s*_ ≡ *c* (*s*), and *r*_*s*_ ≡ *r* (*s*). ***P*** is a matrix with elements *P*_*ss*_′ = *p*_*π*_ (*s*′|*s*). We have defined 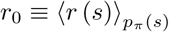 and 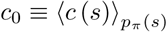 (and thus *L*_*π*_ = *r*_0_ − *λc*_0_). Rearranging:

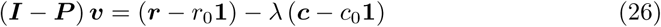

We can solve this system of equations (up to an arbitrary constant, *v*_0_) to find an expression for ***v*** as a linear function of the reward:

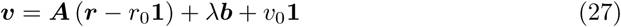

Substituting into Eqn 20, we can express the agent’s policy directly as a function of the reward:

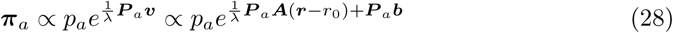

where ***π***_*a*_ is a vector with elements, (***π***_*a*_)_*s*_ ≡ *π* (*a*|*s*) and ***P***_*a*_ is a matrix, with elements (***P*** _*a*_)_*ss*′_ ≡ *p* (*s*′ |*a, s*).

To infer the reward function, *r* (*s*) (up to an irrelevant constant and multiplicative factor, *λ*), we use the observed policy, *π* (*a*|*s*) and transition probabilities, *p* (*s*′|*a, s*), to estimate ***b, A, P***_*a*_ and *p*_*a*_. We then perform numerical optimisation to find the reward that maximises the log-likelihood of the optimal policy in Eqn 28, ⟨log *π*\* (*a*|*s*) ⟩_𝒟_, averaged over observed data, 𝒟.

### Optimising a neural network via RL

We consider a recurrent neural network, with *n* neurons, each described by a binary variable, *σ*_*i*_ = −1*/*1, denoting whether a given neuron is silent/fires a spike in each temporal window. The network receives an external input, ***x***. The network state is described by a vector of *n* binary values, ***σ*** = (*σ*_1_, *σ*_2_, …, *σ*_*n*_)^*T*^. Both the network and input are assumed to have Markov dynamics. Neurons are updated asynchronously, by updating a random neuron at each time-step with probability 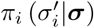. The network dynamics are thus described by:

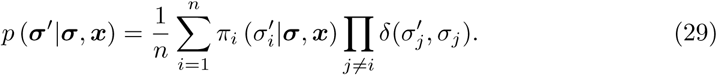

where 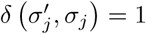 if 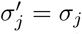 and 0 otherwise.

Equivalently, we can say that at each time, a set of proposed updates, 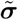, are independently sampled from 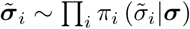, and then a neuron *i* is selected at random to be updated, such that 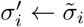.

We define a reward function, *r* (***σ, x***), describing which states are ‘desirable’ for the network to perform a given function. The network dynamics are said to be optimal if they maximise the average reward, ⟨*r* (***σ, x***)⟩_*p*(***σ***,***x***)_ given a constraint on how much each neuron encodes about its inputs 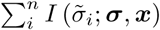. This corresponds to maximising the objective function:

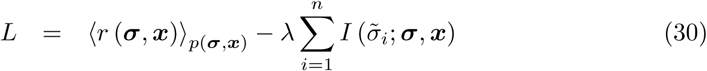

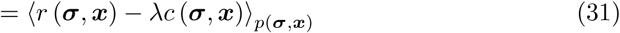

where 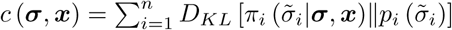 is the coding cost associated with each state, and penalises deviations from each neuron’s average firing rate.

We can decompose the transition probability for the network (Eqn 29), into the probability that a given neuron proposes an update, 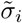, given the network state, ***σ***, and the probability of the new network state, ***σ***′, given 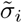 and ***σ***:

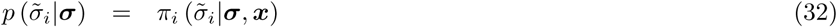

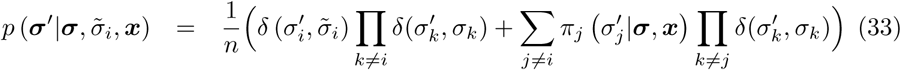

Thus, the problem faced by each neuron, of optimising 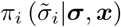 so as to maximise *L*, is equivalent to the MDP described in Methods section, where the action *a*, and state *s* correspond to the neuron’s proposed update 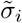 and state of the network and external inputs {***σ, x***}. Thus, we can follow the exact same steps as in Methods section, to show that 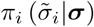 is optimised via the following updates:

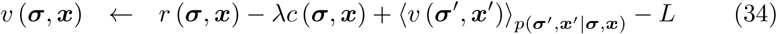

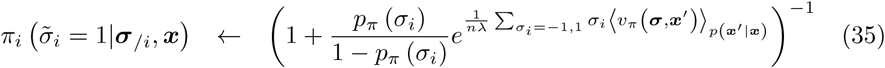

where ***σ***_*/i*_ denotes the state of all neurons except for neuron *i*. As updating the policy for any given neuron increases the objective function, *L*, we can alternate updates for different neurons to optimise the dynamics of the network.

### Inferring network function via inverse RL

After convergence, we can substitute the expression for the optimal policy into the Bellman equality, to obtain:

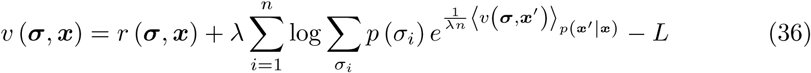

Rearranging, we have:

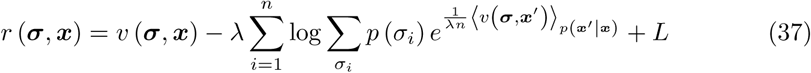

Thus, if we can infer the value function from the observed neural responses, then we can recover the associated reward function through Eqn 37.

To derive an expression for the reward, we first consider the case where there is no external input. In this case, the optimal neural dynamics (Eqn 35) correspond to Gibbs sampling from:

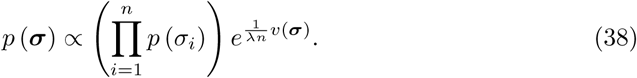

Rearranging, we have a closed-form expression for the value function, *v* (***σ***), in terms of the steady-state distribution:

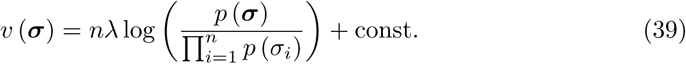

We can then combine Eqn 39 and Eqn 37 to obtain a closed-form expression for the reward function (up to an irrelevant constant):

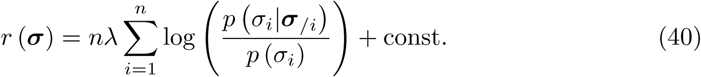

Since we don’t know the true value of *λ*, we can simply set it to unity. In this case, our inferred reward will differ from the true reward by a factor of 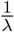. However, since dividing both the reward and coding cost by the same factor has no effect on the shape of the objective function, *L* (but only alters its magnitude), this will not effect any predictions we make using the inferred reward.

With an external input, there is no closed-form solution for the value function. Instead, we can infer *v* (***σ, x***) numerically by maximising the log-likelihood, 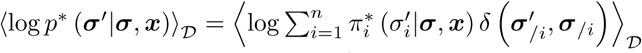, where 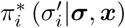 denote optimal response probabilities. Once we know *v* (***σ, x***) we can compute the reward from Eqn 37.

### Approximate method for larger networks

#### RL model

To scale our framework to larger networks we approximate the value function, *v* (***σ, x***), by a parametric function of the network activity, ***σ*** and input, ***x***. Without loss of generality, we can parameterise the value function as a linear combination of basis functions: *v*_***ϕ***_ (***σ, x***) ≡ ***ϕ***^*T*^ ***f*** (***σ, x***). From Eqn 36, if the network is optimal, then the value function equals:

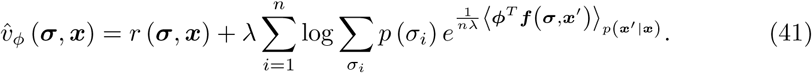

In the exact algorithm, we updated the value function by setting it equal to 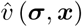 (Eqn 34). Since in the parametric case this is not possible, we can instead update ***ϕ*** to minimise:

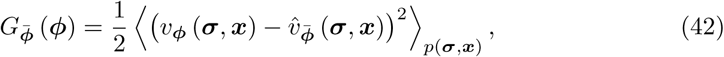

where 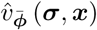 is the target value function, defined as in Eqn 41, with parameters, 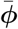.

We follow the procedure set out in [16, 17] to transform this into a stochastic gradient descent algorithm. First, we perform *n*_*batch*_ samples from the current policy. Next, we perform a stochastic gradient descent update:

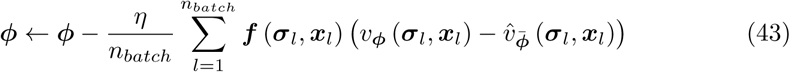

where *η* is a constant that determines the learning rate. Finally, after doing this *n*_*epoch*_ times, we update the target parameters, 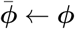. These steps are repeated until convergence.

#### Inverse RL

We can infer the parameters of the value function, ***ϕ***, by maximising the log-likelihood: ⟨log *p*_***ϕ***_ (***σ***′|***σ, x***)⟩_𝒟_. We can choose the form of the value function to ensure that this is tractable. For example, if the value function is quadratic in the responses, then this corresponds to inferring the parameters of a pairwise Ising model [27, 28].

After inferring ***ϕ***, we want to infer the reward function. At convergence, ∇_*ϕ*_ *F* (*ϕ*) = 0 and 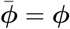, so that:

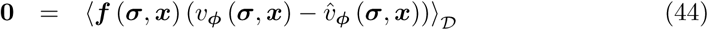

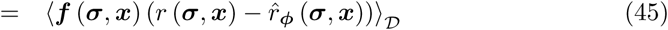

where

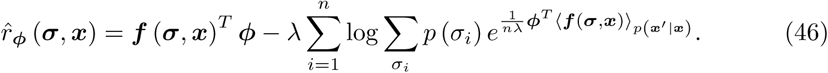

In the exact case, where 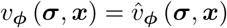 (and thus, 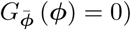), then the inferred reward equals 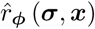. However, this is not necessarily true when we assume an approximate value function.

Just as we did for the value function, we can express the reward function as a linear combination of basis functions: *r* (***σ, x***) = ***θ***^*T*^ ***g*** (***σ, x***). Thus, Eqn 45 becomes:

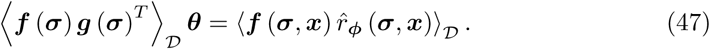

If the reward function has the same number of parameters than the approximate value function (i.e. ***f*** (***σ***) and ***g*** (***σ***) have the same size), then we can solve this equation to find ***θ***. Alternatively, if the reward function has more parameters than the value function, then we require additional assumptions to unambiguously infer the reward.

### Simulation details

#### Agent navigating a maze

We considered an agent navigating a 15 × 15 maze. The agent’s state corresponded to their position in the maze. The agent could choose to move up, down, left or right in the maze. At each time there was a 5% probability that the agent moved in a random direction, independent of their selected action. Moving in the direction of a barrier (shown in blue in Fig 1D) would result in the agent remaining in the same location. After reaching the ‘rewarded’ location (bottom right of the maze), the agent was immediately transported to a starting location in the top left of the maze. We optimised the agent’s policy at both low and high coding cost (*λ* = 0.013*/*0.13 respectively) using the entropy-regularised RL algorithm described in Methods section. The reward was inferred from the agent’s policy after optimisation as described in Methods section.

#### Network with single binary input

We simulated a network of 8 neurons that receive a single binary input, *x*. The stimulus has a transition probability: *p* (*x*′ = 1|*x* = −1) = *p* (*x*′ = −1|*x* = 1) = 0.02. The reward function was unity when *x* = −1 and the network fired exactly 2 spikes, or when *x* = −1 and the network fired exactly 6 spikes. We set *λ* = 0.114.

To avoid trivial solutions where a subset of neurons spike continuously while other neurons are silent, we defined the coding cost to penalise deviations from the population averaged firing rate (rather than the average firing rate for each neuron). Thus, the coding cost was defined as 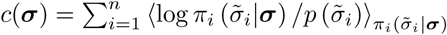, where 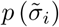 is the average spiking probability, across all neurons.

We inferred the reward *r* (***σ***) from neural responses as described in Methods section. Note that, when inferring the reward, it did not matter if we assumed that the constraint included the population averaged firing rate or the average firing rate for each neuron individually, since after optimisation all neurons had the same mean firing rate. In Fig 2E we rescaled and shifted the inferred reward to have the same mean and variance as the true reward.

We used the inferred reward to predict how neural tuning curves should adapt when we alter the stimulus statistics (Fig 2F, upper) or remove a cell (Fig 2F, lower). For fig 2F (upper), we altered the stimulus statistics by setting *p* (*x*′ = 1|*x* = −1) = 0.01 and *p* (*x*′ = −1|*x* = 1) = 0.03. For figure 2C (lower), we removed one cell from the network. In both cases, we manually adjusted *λ* to keep the average coding cost constant.

#### Efficient coding

We considered a stimulus consisting of *m* = 7 binary variables, *x*_*i*_ = −1*/*1. The stimulus had Markov dynamics, with each unit updated asynchronously. The stimulus dynamics were given by:

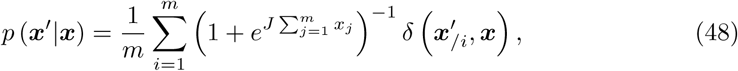

where *J* = 1.5 is a coupling constant. A ‘relevant’ variable, *y* (***x***) was equal to 1 if 4 or more inputs equalled 1, and equal to −1 otherwise.

We optimised a network of *n* = 7 neurons to efficiently code the relevant variable *y* (***x***), using the algorithm described in the main text. For Fig 4B-C we set *λ* = 0.167. For Fig 4D we varied *λ* between 0.1 and 0.5. For Fig 4E we altered the stimulus statistics so that,

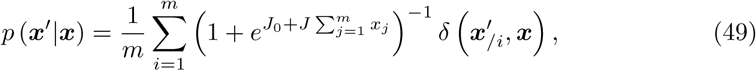

where *J*_0_ was a bias term that we varied between 0 and 0.4. For each value of *J*_0_ we adjust *λ* so as to keep the average coding cost constant.

#### Pairwise coupled network

We considered a network of 12 neurons arranged in a ring. We defined a reward function that was equal to 1 if exactly 4 adjacent neurons were active, and 0 otherwise. We defined the coding cost as described in the previous section, to penalise deviations from the population averaged mean firing rate.

We approximated the value function by a quadratic function, *v* (***σ***) = ∑ _*i,j*≠*i*_*J*_*ij*_*σ*_*i*_*σ*_*j*_ + ∑ _*i*_*h*_*i*_*σ*_*i*_. We optimised the parameters of this value function using the algorithm described in Methods section, with *λ* = 0.05. We used batches of *n*_*batch*_ = 40 samples, and updated the target parameters after every *n*_*epoch*_ = 100 batches.

We inferred the reward function from the inferred network couplings, ***J*** and ***h***. As described in Methods section, this problem is only well-posed if we assume a low-d parametric form for the reward function, or add additional assumptions. We therefore considered several different sets of assumptions. For our initial ‘sparse model’, we set up a linear programming problem in which we minimised *l*_1_ = ∑_***σ***_ *r* (***σ***), under the constraint that the reward was always greater than 0 while satisfying the optimality criterion given by Eqn 47. For the ‘pairwise model’ we assumed that *r* = ∑_*i,j*_*W*_*ij*_*σ*_*i*_*σ*_*j*_. We fitted the parameters, *W*_*ij*_, so as to minimise the squared difference between the left and right hand side of Eqn Eqn 47. Finally, for the ‘global model’ we assumed that *r* =∑_*j*_ *δ*_*j,m*_*W*_*j*_, where *m* is the total number of active neurons and *δ*_*ij*_ is the kronecker-delta. Parameters, *W*_*j*_ were fitted to the data as for the pairwise model.

Finally, for the simulations shown in Figure 5, panels B-D, we ran the optimisation with *λ* = 0.1, 0.01 and 0.1, respectively. For panel 3C we removed connections between neurons separated by a distance of 3 or more on the ring. For panel 3D we forced two of the neurons to be continuously active.

## Acknowledgements

This work was supported by ANR JCJC grant (ANR-17-CE37-0013) to M.C, ANR Trajectory (ANR-15-CE37-0011), ANR DECORE (ANR-18-CE37-0011), the French State program Investissements d’Avenir managed by the Agence Nationale de la Recherche (LIFESENSES; ANR-10-LABX-65), EC Grant No. H2020-785907 from the Human Brain Project (SGA2), and an AVIESAN-UNADEV grant to O.M. The authors would like to thank Ulisse Ferrari for useful discussions and feedback.

## Notes

#### Summary of Updates

We have changed the title. We have added two additional figures (fig 2 and fig 6). We have made changes to the text to highlight the limitations/scope of our approach.

## References

1. Yang G R, Joglekar MR, Song HF, Newsome WT, Wang XJ. (2019). Task representations in neural networks trained to perform many cognitive tasks. Nat Neurosci, 22:297–306s

2. Heeger D J (2017) Theory of cortical function. Proc Natl Acad Sci USA 114:1773–1782

3. Sussillo D, Abbott L F (2009) Generating coherent patterns of activity from chaotic neural networks. Neuron 63:544–557.

4. Gütig R (2016) Spiking neurons can discover predictive features by aggregate–label learning. Science 351(6277):aab4113

5. Hopfield JJ (1982) Neural networks and physical systems with emergent collective computational abilities Proc Natl Acad Sci USA 79:2554–2558

6. Körding K (2007) Decision theory: what should the nervous system do? Science 318:606–610

7. Boerlin M, Machens CK, Denève S (2013) Predictive coding of dynamical variables in balanced spiking networks. PLoS Comp Bio 9 e1003258.

8. Simoncelli EP, Olshausen BA (2001) Natural image statistics and neural representation. Ann Rev Neurosci 24:1193–1216

9. Tkačik G, Prentice JS, Balasubramanian V, Schneidman E (2010) Optimal population coding by noisy spiking neurons. Proc Natl Acad Sci USA 107:14419–14424.

10. Chalk M, Marre O, Tkačik G (2018) Toward a unified theory of efficient, predictive, and sparse coding. Proc Natl Acad Sci USA 115:186–191.

11. Barlow, HB (1961) Possible principles underlying the transformations of sensory messages. Sensory Communication, ed Rosenblith WA (MIT Press, Cambridge, MA), pp 217–234

12. Field DJ (1994) What is the goal of sensory coding? Neural Comput 6:559–601.

13. Gjorgjieva J, Sompolinsky H, Meister M (2014) Benefits of pathway splitting in sensory coding. J Neurosci 34:12127–12144.

14. Sutton RS, Barto AG (2018) Reinforcement learning: An introduction. MIT press.

15. Todorov E (2008) General duality between optimal control and estimation. Proc of the 47th IEEE Conference on Decision and Control 4286–4292

16. Schulman J, Chen X, Abbeel P (2017) Equivalence between policy gradients and soft Q-learning. arXiv:1704.06440

17. Haarnoja T, Tang H, Abbeel P, Levine S (2017). Reinforcement learning with deep energy-based policies. Proc 34th International Conf on Machine Learning 70:1352–1361

18. Tiomkin S, Tishby N (2017). A Unified Bellman Equation for Causal Information and Value in Markov Decision Processes. arXiv:1703.01585.

19. Mahadevan S. (1996). Average reward reinforcement learning: Foundations, algorithms, and empirical results. Machine learning 22:159–195.

20. Ng AY, Russell SJ (2000) Algorithms for inverse reinforcement learning. Proc of the 17th International Con on Machine Learning pp. 663–670

21. Rothkopf CA, Dimitrakakis C (2011) Preference elicitation and inverse reinforcement learning. In Joint European conference on machine learning and knowledge discovery in databases Springer pp. 34–48.

22. Herman M, Gindele T, Wagner J, Schmitt F, Burgard W (2016) Inverse reinforcement learning with simultaneous estimation of rewards and dynamics. Artificial Intelligence and Statistics 102–110

23. Wu Z, Schrater P, Pitkow X (2018) Inverse POMDP: Inferring What You Think from What You Do. arXiv:1805.09864.

24. Reddy S, Dragan AD, Levine S (2018) Where Do You Think You’re Going?: Inferring Beliefs about Dynamics from Behavior. arXiv:1805.08010.

25. Berger T. Rate Distortion Theory. (1971) Englewood Clis.

26. Bialek W, van Steveninck RRDR, Tishby N (2006) Efficient representation as a design principle for neural coding and computation. IEEE international symposium on information theory 659–663

27. Schneidman E, Berry MJ, Segev R, Bialek W (2006) Weak pairwise correlations imply strongly correlated network states in a neural population. Nature 440:1007–1012

28. Tkačik G, Marre O, Amodei D, Schneidman E, Bialek W, Berry MJ (2014) Searching for collective behavior in a large network of sensory neurons. PLoS Comp Bio 10:e1003408.

29. Ben-Yishai R, Bar-Or RL, Sompolinsky H (1995) Theory of orientation tuning in visual cortex. Proc Natl Acad Sci, 92:3844–3848

30. Zhang K (1996) Representation of spatial orientation by the intrinsic dynamics of the head-direction cell ensemble: a theory. J Neurosci 16:2112–2126.

31. Kim SS, Rouault H, Druckmann S, Jayaraman V (2017) Ring attractor dynamics in the Drosophila central brain. Science 356:849–853.

32. Pillow JW, Shlens J, Paninski L, Sher A, Litke AM, Chichilnisky EJ, Simoncelli EP (2008) Spatio-temporal correlations and visual signalling in a complete neuronal population. Nature 454:995–999

33. McIntosh L, Maheswaranathan N, Nayebi A, Ganguli S, Baccus S (2016) Deep learning models of the retinal response to natural scenes. Adv Neur Inf Proc Sys 29:1369–1377

34. Cunningham JP, Yu BM (2014) Dimensionality reduction for large-scale neural recordings. Nat Neurosci 17:1500–1509

35. Rubin A, Sheintuch L, Brande-Eilat N, Pinchasof O, Rechavi Y, Geva N, Ziv Y (2019) Revealing neural correlates of behavior without behavioral measurements. bioRxiv:540195

36. Chaudhuri R, Gercek B, Pandey B, Peyrache A, Fiete I (2019) The population dynamics of a canonical cognitive circuit. bioRxiv: 516021

37. Goddard E, Klein C, Solomon SG, Hogendoorn H, Carlson TA (2018) Interpreting the dimensions of neural feature representations revealed by dimensionality reduction NeuroImage 180:41–67

38. Sharpee T, Rust NT, Bialek W (2003) Maximally informative dimensions: analyzing neural responses to natural signals. Adv Neur Inf Proc Sys 277–284

39. Niv Y (2009) Reinforcement learning in the brain. J Mathemat Psychol 53:139–154

40. Dayan P, Niv Y (2008) Reinforcement learning: the good, the bad and the ugly. Curr Op Neurobio 18:185–196.

41. Daw ND, Doya K (2006) The computational neurobiology of learning and reward. Curr Op Neurobio 16:199–204.

42. Fairhall AL, Geoffrey DL, William B, de Ruyter van Steveninck RR. (2001) Efficiency and ambiguity in an adaptive neural code. Nature 412:787.

43. Benucci A, Saleem AB, Carandini M. (2013). Adaptation maintains population homeostasis in primary visual cortex. Nat Neurosci 16:724.

44. Li N, Kayvon D, Karel S, and Shaul D. (2016) Robust neuronal dynamics in premotor cortex during motor planning. Nature. 532:459.

45. Mlynarski W, Hledik M, Sokolowski TR, Tkacik G (2019). Statistical analysis and optimality of biological systems. bioRxiv:848374.

46. Aenugu S, Abhishek S, Sasikiran Y, Hananel H, Thomas PS, Kozma R. (2019) Reinforcement learning with spiking coagents. arXiv:1910.06489

